# Resting-state EEG recorded with gel-based versus consumer dry electrodes: spectral characteristics and across-device correlations

**DOI:** 10.1101/2023.08.09.552601

**Authors:** Daria Kleeva, Ivan Ninenko, Mikhail Lebedev

## Abstract

Recordings of electroencephalographic (EEG) rhythms and their analyses have been instrumental in basic Neuroscience, clinical diagnostics, and the field of brain-computer interfaces (BCIs). While in the past such measurements have been conducted mostly in laboratory settings, recent advancements in dry electrode technology pave way to a broader range of consumer and medical application because of their greater convenience compared to gel-based electrodes. Here we conducted resting-state EEG recordings in two groups of healthy participants using three dry-electrode devices, the Neiry Headband, the Neiry Headphones and the Muse Headband, and one standard gel electrode-based system, the NVX. We examined signal quality for various spatial and spectral ranges which are essential for cognitive monitoring and consumer applications. Distinctive characteristics of signal quality were found, with the Neiry Headband showing sensitivity in low-frequency ranges and replicating the modulations of delta, theta and alpha power corresponding to the eyes-open and eyes-closed conditions, and the NVX system performing well in capturing high-frequency oscillations. The Neiry Headphones were more prone to low-frequency artifacts compared to the Neiry Headband, yet recorded modulations in alpha power and had a strong alignment with the NVX at higher frequencies. The Muse Headband had several limitations in signal quality. We suggest that while dry-electrode technology appears to be appropriate for the EEG rhythm-based applications the potential benefits of these technologies in terms of ease of use and accessibility should be carefully weighted against the capacity of each concrete system.

## 1. Introduction

Electroencephalography (EEG) has been an invaluable tool in neuroscientific research, clinical diagnostics, and brain-computer interface (BCI) applications. Conventionally, EEG recordings have relied on wet gel-based electrodes which necessitate conductive gel application, diligent maintenance, wired digital amplifiers, and a direct computer connection for data storage. However, recent advancements in dry electrode technology have offered a promising alternative, reducing the complexities posed by conventional gel electrodes. These novel dry electrode systems are designed to be user-friendly, opening new possibilities for improving the overall user experience and real-world EEG applications outside the restrictions of laboratory settings (Niso et al., 2023).

The quality of EEG signal acquisition is of paramount importance in any application. Signal integrity directly impacts the accuracy and reliability of EEG-based studies, including cognitive state monitoring, emotion recognition, and motor control research, among others. Therefore, rigorous validation and comparison of the signal quality between the emerging dry electrode systems and conventional gel electrodes are crucial to understanding their potential benefits and drawbacks.

Particular emphasis should be placed on evaluating the quality of capturing resting-state EEG, rather than focusing solely on responses evoked by stimuli. This distinction is crucial for consumer EEG applications that center around cognitive monitoring. Resting-state EEG data, owing to its capacity for signal processing and analysis, can unveil intricate patterns tied to attention, memory, and emotional states (Katahira et al., 2018; Bitner and Le, 2022; Sun and Yeh, 2017; Bajada and Bonello, 2021; Trejo et al., 2015; Myrden and Chau, 2017). This, in turn, may allow users to track their cognitive well-being over time, optimize their daily routines, and make well-informed decisions regarding lifestyle choices that have the potential to impact cognitive performance.

In this study, we conducted resting-state EEG recordings using three modern dry-electrode devices: the Neiry Headband and Neiry Headphones by Neiry LLC, Russia, and the Muse S Headband by InterAxon LLC, Canada. Additional consumer devices will be evaluated with the same method upon availability. The Neiry Headband is a novel commercially available device developed for working with EEG-based paradigms based on EEG spectral characteristics. There have been no previous studies validating the signal quality of the Neiry devices or using them for other research purposes.

The third device that was available to us, the Muse Headband, is a widely used commercial device intended for aiding meditation and sleep. Several studies have investigated its signal quality. These studies have shown that, on the one hand, it could successfully capture typical EEG spectral characteristics, such as the alpha spectral peak. However, on the other hand, it exhibited low reliability in terms of repeatability of measures conducted on different days (Ratti et al., 2017). Additionally, the Muse Headband struggled to reproduce typical brain oscillations and was susceptible to the influence of artifacts (Przegalinska et al., 2018). These reported drawbacks of the Muse Headband contributed to our conservative expectations for the Neiry Headband, as well.

Despite certain limitations compared to the medical-grade systems, the Muse Headband has demonstrated its effectiveness for ERP research (Krigolson et al., 2017). Additionally this device has found applications in various areas, including predicting task performance (Papakostas et al., 2017), human stress classification (Asif et al., 2019), emotion recognition (Bano et al., 2022), perception of mental stress (Arsalan et al., 2019), and other purposes.

Consequently, we opted to use the Muse Headband as a point of comparison to assess whether the signals obtained from the Neiry Headband or Neiry Headphones exhibited similar properties. To serve as a control device representative traditional EEG recordings, we selected the NVX EEG System (MCS, Russia) with gel electrodes, as it fulfilled the necessary criteria for a standard medical EEG recording device.

Consistent with the previous validation studies conducted with the use of other devices (Przegalinska et al., 2018; Ratti et al., 2017; Wyckoff et al., 2015; Cannard et al., 2021; Marini et al., 2019; Kam et al., 2019; Radüntz, 2018), the present study established three measures of signal quality. First, the increase in alpha power was examined during eyes-closed compared to eyes-open conditions. Second, the power correspondence in the standard frequency bands was assessed in comparison to the gel apparatus. Finally, the correlation of measures of spectral power between the devices was investigated.

## 2. Methods

### 2.1. Participants

Two distinct research studies were conducted to compare the Neiry Headband and Neiry Headphones with NVX and Muse Headband. Each study enlisted 15 healthy volunteers initially. Following the removal of participants whose EEG data exhibited substantial artifacts caused by power-line interference, the first study ultimately comprised eleven healthy volunteers (consisting of five males and six females, with an average age of 26.3 years), while the second study included thirteen volunteers (three males and ten females with an average age of 22.8 years). The participants had normal or corrected-to-normal vision and provided written informed consent, following the ethics protocol approved by the Ethics Committee of Skolkovo Institute of Science and Technology (Protocol No.9 of Institutional Review Board, June 22, 2022).

### 2.2. Data acquisition

During the experiment, participants were seated comfortably and asked to undergo two phases of EEG recording. The first phase involved a five-minute session with eyes open, followed by an additional five-minute session with eyes closed.

The resting-state recordings were obtained using three different devices in the following sequence: Neiry Headband or Neiry Headphones (Neiry LLC, Russia), Muse S Headband (InterAxon LLC, Canada), and NVX-36 amplifier with gel-based Ag/Cl wired electrodes (Medical Computer Systems Ltd., Russia). The Neiry Headband is a soft band placed around the head, equipped with four dry EEG electrodes (T3, T4, O1, O2), each containing 25 pogo pins (spring-loaded electric connectors). The reference and ground electrodes are positioned frontally. The Neiry Headphones come with four dry EEG electrodes, specifically positioned at C3, C4, TP9, and TP10. The Muse system uses electrodes located similarly to Fpz, AF7, AF8, TP9, and TP10, with electrode Fpz serving as the reference electrode. In the NVX system, the electrode locations were comparable to those in the Neiry Headband (T3, T4, O1, O2), or Neiry Headphones (C3, C4, TP9, TP10) and Fp1-Fp2 electrodes were used for referencing, while ground electrode was placed as Fpz. The choice of reference and ground electrodes in NVX closely replicated the setup in Neiry Headband.

During the recordings, the default sampling rates for the portable devices were as follows: 250 Hz for the Neiry Headband, 256 Hz for the Muse Headband, and 250 Hz for the NVX system. To ensure optimal signal quality in the recordings, electrode impedance was checked before starting the EEG recording, ensuring that it was below 50 kΩ for NVX and below 500 kΩ for Neiry Headband. As for the Muse Headband, direct access to impedance values was not possible, but the available indicators showed that the impedance level was satisfactory.

### 2.3. EEG processing

The data analyses were performed using MNE Python software (Gramfort et al., 2013) along with other Python libraries. After importing the recorded data into MNE Python, the Muse recordings were resampled at 250 Hz. Subsequently, the data underwent high-pass filtering from 0.5 Hz using an overlap-add Finite Impulse Response (FIR) filter. To conduct spectral analysis on the time series from each channel, we computed the Power Spectral Density (PSD) using Welch’s method (Welch, 1967).

Next, we extracted the mean logarithm of PSD (log PSD) values for several frequency bands (delta: 1.5-3.5 Hz, theta: 4–7.5 Hz, alpha: 8–12 Hz, low beta: 13–16 Hz, beta: 13–21 Hz, high beta: 21–32 Hz, gamma: 32-40 Hz) at each electrode site for further analysis.

### 2.4. Statistical analysis

The statistical analyses employed in this study utilized a repeated-measures, within-subject Analysis Of Variance (ANOVA) with participants considered as a random variable and *Device, Condition* and *Channel group* treated as within-subject factors. Since the portable devices allow to record activity only from four sensors, we did not examine signal quality in terms of spatial resolution and merged the features within the anterior (T3 and T4 for Neiry Heabdand and NVX, AF7 and AF8 for Muse) and posterior sites (O1 and O2 for Neiry Headband and NVX, TP9 and TP10 for Muse). To conduct post-hoc pairwise comparisons, t-tests were utilized, and a Bonferroni correction was implemented to account for multiple comparisons.

## 3. Results

### 3.1. Study 1. Neiry Headband

#### 3.1.1. Resting-state EEG

In Figure 1, 10-s samples of resting-state EEG data are depicted for one subject for the eyes-open and eyes-closed conditions. Data acquired from three different devices are shown. Upon a visual examination, the presence of alpha spindles is clear during the eyes-closed condition for all three devices. However, for the Muse recordings (Fig. 1, 3.b), these alpha spindles have a lower amplitude compared to the other devices.

**Figure 1:**
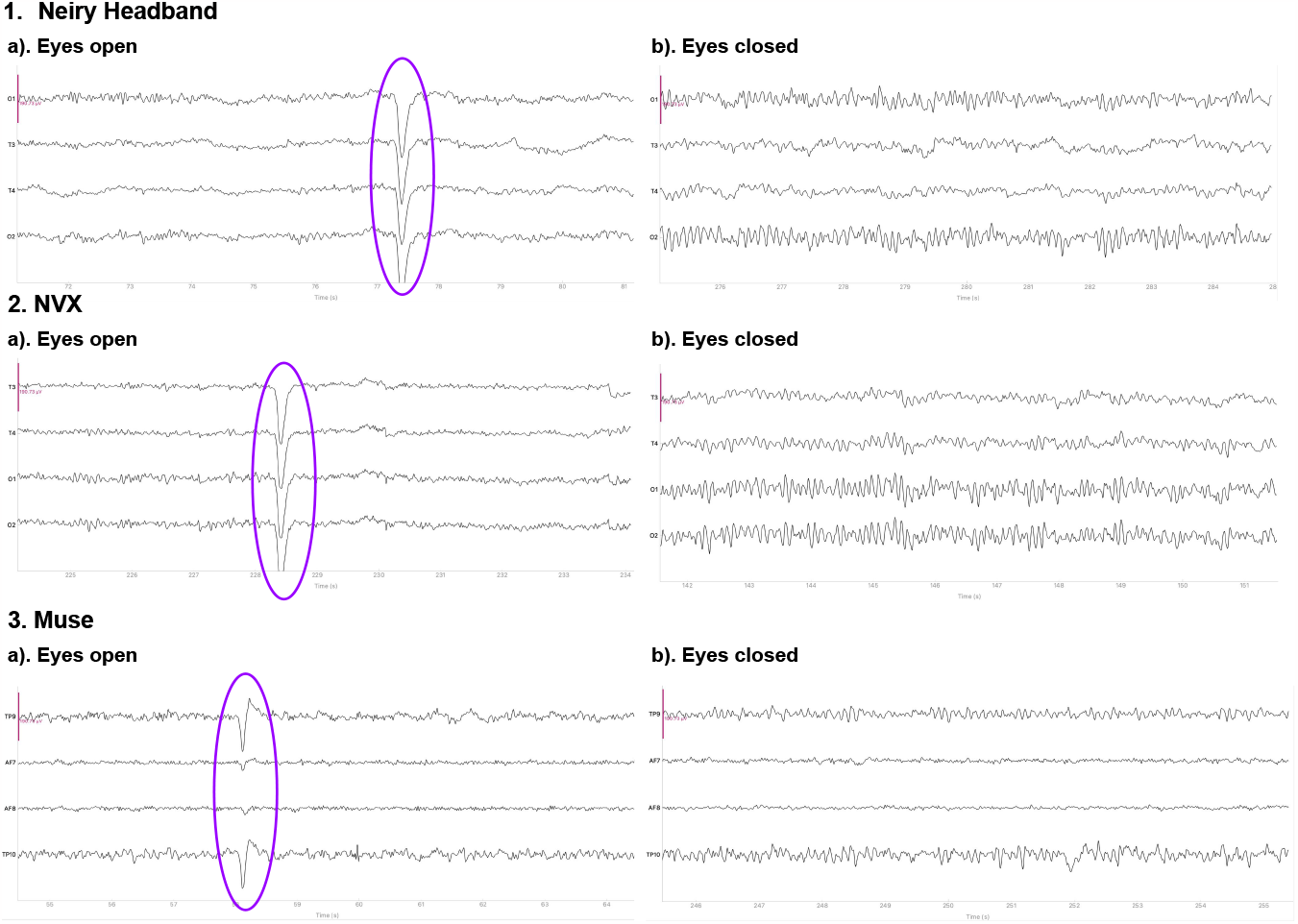
Representative samples of the recordings with the Neiry Headband (1), NVX (2) and Muse (3) devices for the eyes-open and eyes-closed conditions. The purple ovals mark the typical eye-blink artifacts. The y-limit is set to ±190.73 mV for all graphs. The signal was bandpass-filtered from 0.5 to 30 Hz for visualization.

**Figure 2:**
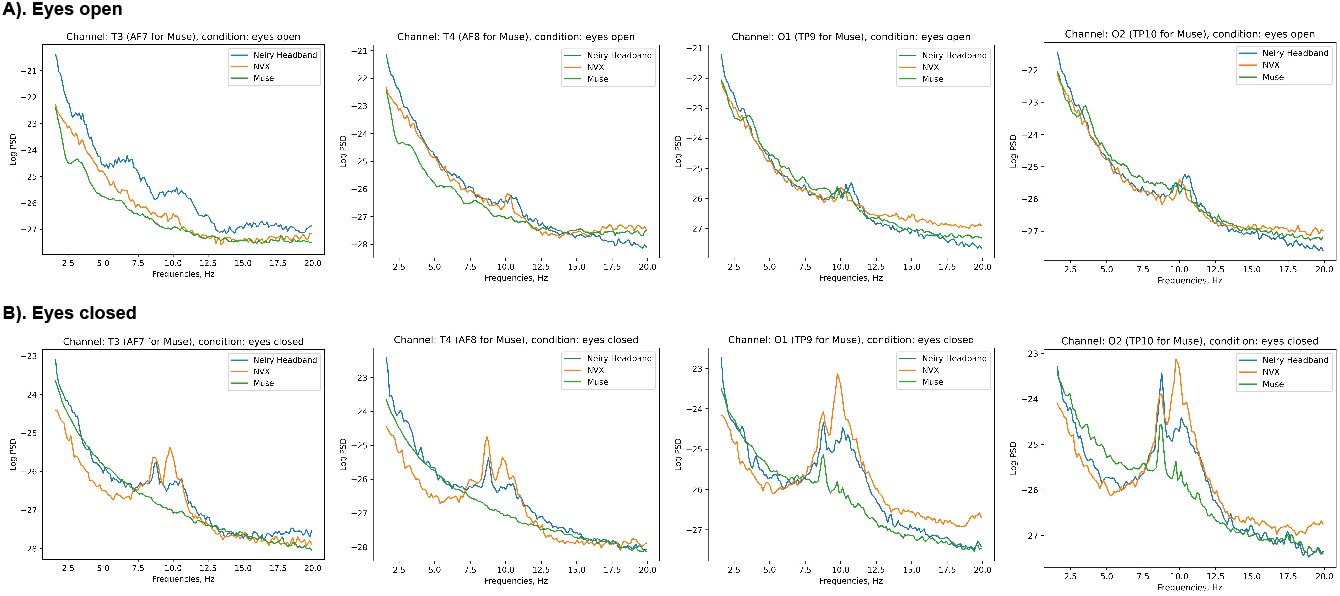
Across-subject (N=11) averages of log PSD demonstrating the difference in alpha power during eyes-open (A) versus eyes-closed (B) conditions for Neiry Headband, NVX and Muse devices

**Figure 3:**
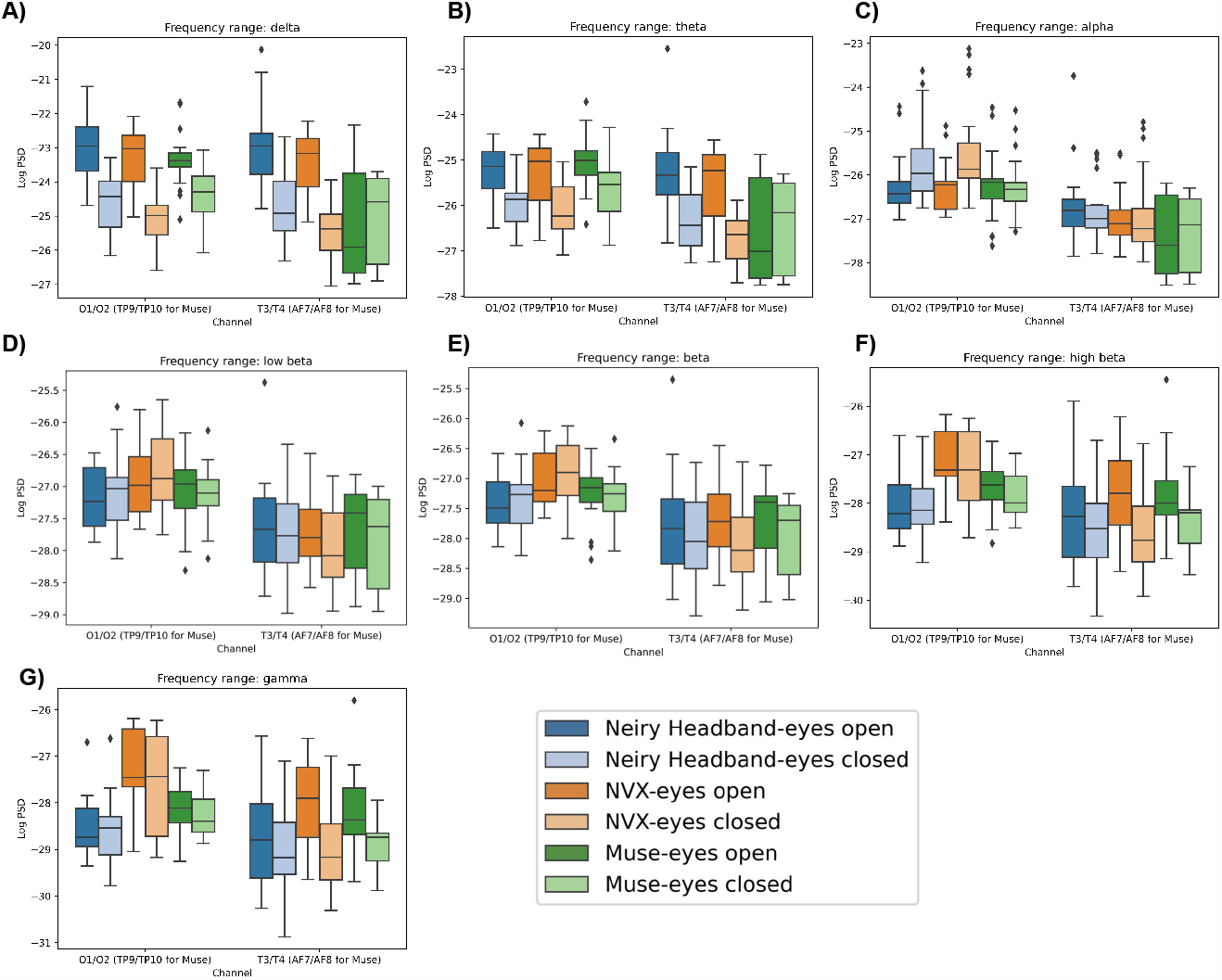
Boxplots showing the distribution of mean log PSD values across conditions and channel groups for different frequency ranges: delta (A), theta (B), alpha(C), low beta (D), beta (E), high beta (F), gamma (G) for Neiry Headband, NVX and Muse

#### 3.1.2. Spectral analysis

Figure 2 shows the across-participant average of log PSD, clearly showing an increase in alpha power when the eyes were closed.

A repeated measures ANOVA was conducted to examine the effects of device, condition and channel group on the mean value of log PSD in a given frequency range.

The results showed that the choice of a device did not have a significant main effect on the delta power (F(2, 20) = 2.611, p = 0.0983) (Fig. 3, A). However, both condition (F(1, 10) = 63.3793, p < 0.001) and channel group (F(1, 10) = 27.2044, p < 0.001) had significant main effects. Moreover, there were significant interactions observed between the device and condition (F(2, 20) = 5.3061, p < 0.05), device and channel group (F(2, 20) = 16.3133, p < 0.001), condition and channel group (F(1, 10) = 8.9950, p < 0.05), as well as device, condition, and channel group (F(2, 20) = 9.6415, p < 0.05).

The post-hoc analysis revealed that when participants had their eyes closed, delta power was significantly reduced compared to the eyes open conditions (t=-11.853, p<0.001). Additionally, delta power was stronger over the posterior group of channels compared to the anterior group (t=5.9207, p<0.001). In the eyes open condition, recordings from Neiry Headband exhibited higher delta power compared to the NVX recordings (t=2.811, p<0.05) and Muse recordings (t=4.602, p<0.001). Moreover, NVX recordings had higher delta power than Muse recordings (t=3.350, p<0.05). Conversely, during the eyes closed condition, delta power was stronger in the recordings from Neiry Headband compared to NVX recordings (t=4.345, p<0.001). However, NVX recordings had lower delta power than Muse recordings (t=-2.407, p<0.05). Notably, delta power substantially increased after the eyes opened for both the Neiry Headband recordings (t=12.339, p<0.001) and NVX recordings (t=17.397, p<0.001).

The delta power was significantly different across the devices and sites. Specifically, the Neiry Headband recordings consistently yielded higher delta power compared to NVX recordings over both the posterior (t=2.915, p<0.05) and anterior sites (t=4.252, p<0.001). In comparison to Muse, Neiry Headband recordings had higher delta power only for the anterior sites (t=4.158, p<0.001). Notably, there was a significant decrease in delta power over the anterior sites compared to posterior sites, but this pattern was observed only for the NVX (t=-10.932, p<0.001) and Muse recordings (t=-6.717, p<0.001).

For the eyes-open condition considered separately, delta power over the anterior sites was stronger for the Neiry Headband recordings as compared to Muse (t=5.049, p<0.001), and in the NVX recordings as compared to Muse (t=3.950, p<0.05). For the eyes-closed condition, delta power was stronger for the Neiry Headband than for NVX for both the anterior (t=3.582, p<0.05) and posterior (t=2.511, p<0.05) sites. Furthermore, a statistically significant positive difference in delta power between the eyes-open and eyes-closed conditions was found for the Neiry recordings over the posterior (t=8.828, p<0.001) and anterior sites (t=8.454, p<0.001), for NVX over the posterior sites (t=11.368, p<0.001) and anterior sites (t=13.206, p<0.001), and for Muse only over the posterior sites (t=4.484, p<0.05).

The ANOVA analysis conducted for the theta band data (Fig. 3, B) revealed that there was no significant main effect of the recording device (F(2,20) = 0.6968, p = 0.5099). However, both condition (F(1,10) = 16.2164, p < 0.05) and channel group (F(1,10) = 99.6514, p < 0.001) had significant main effects. Interactions between device and condition (F(2,20) = 2.9222, p = 0.0770) and condition and channel (F(1,10) = 0.1589, p = 0.6986) were found to be non-significant. On the other hand, the interactions between device and channel (F(2,20) = 12.9638, p < 0.001) and device, condition, and channel (F(2,20) = 10.0947, p < 0.001) were statistically significant.

When participants had their eyes closed, theta power was significantly suppressed (t = -8.5827, p < 0.001). Furthermore, theta power was more prominent over the posterior sites compared to the anterior sites (t = 9.4043, p < 0.001).

The theta power exhibited distinct patterns across the different recording devices. Specifically, in the Neiry Headband recordings, theta power was lower compared to Muse recordings over the posterior sites (t = -2.77, p < 0.05), while it was larger at the anterior sites (t = 2.875, p < 0.05). On the other hand, in NVX recordings, theta power was suppressed over the posterior sites compared to Muse (t = -3.656, p < 0.05) and over the anterior sites compared to Neiry Headband (t = -3.718, p < 0.05). Furthermore, a general tendency for the theta power being lower at the anterior sites compared to the posterior ones was observed for both NVX (t = -10.471, p < 0.001) and Muse recordings (t = -8.9333, p < 0.001).

The theta power had significant differences across the recording devices and eye conditions. When the eyes were open, Neiry Headband’s theta power was significantly higher compared to Muse for the anterior regions (t=3.495, p<0.05). Conversely, the NVX’s theta power was significantly lower compared to Muse for the posterior regions when the eyes were closed (t=-3.17, p<0.05). Additionally, the theta power in the Neiry Headband recordings was significantly higher than in the NVX recordings for the anterior regions when the eyes were closed (t=3.387, p<0.05).

Furthermore, when the eyes were open, the theta power over the posterior regions increased for the Neiry Headband (t=5.535, p<0.001), NVX (t=5.785, p<0.001) and Muse (t=3.633, p<0.05). Over the anterior channels, this increase was observed only for the Neiry Headband (t=6.313, p<0.001) and NVX (t=8.033, p<0.001).

The ANOVA conducted for the alpha frequency range (Fig. 3, C) yielded significant main effects of the recording device (F(2,20)=3.9126, p<0.05) and channel group (F(1,10)=155.2894, p<0.001). However, the main effect of condition did not reach significance (F(1,10)=1.8573, p=0.2028). The interactions between device and condition (F(2,20)=1.1401, p=0.3397) and device and channel (F(2,20)=2.2727, p=0.1290) were also non-significant. In contrast, the interactions between condition and channel group were significant (F(1,10)=18.1444, p<0.05), as well as the interactions between device, condition, and channel group (F(2,20)=9.3815, p<0.05).

For the posterior regions, the alpha power was significantly stronger when the eyes were closed compared to when they were open (t=-4.295, p<0.001). This effect was statistically significant for the Neiry Headband (t=-3.199, p<0.05) and NVX (t=-4.739, p<0.001) recordings. Additionally, when the eyes were closed, the Neiry Headband recordings had higher alpha power for the posterior channels than the recordings with Muse (t=5.366, p<0.001), and NVX had higher alpha power for the posterior channels compared to Muse (t=4.855, p<0.001).

The ANOVA analysis conducted for the low beta frequency range (Fig. 3, D) showed a significant main effect of the channel group (F(1,10) = 85.0479, p < 0.001). The main effects of the recording device (F(2,20) = 0.3708, p = 0.6949) and experimental condition (F(1,10) = 0.4008, p = 0.5409) were not statistically significant. The interaction between device and condition (F(2,20) = 0.0467, p = 0.9544) and the interaction between device, condition, and channel group (F(2,20) = 2.3409, p = 0.1220) were not significant either. The only significant interactions were found between device and channel (F(2,20) = 4.7482, p < 0.05) and between condition and channel (F(1,10) = 15.2382, p < 0.05).

The low-beta power was stronger for the posterior than anterior channels (t = 16.0988, p < 0.001). Additionally, NVX recordings exhibited stronger low-beta power for the posterior channels as compared to both Neiry Headband (t = 4.9032, p < 0.001) and Muse (t = 2.353, p < 0.05).

Similar to the findings for the low beta power, the ANOVA analysis conducted on beta power (Fig. 3, E) also revealed a significant main effect of the channel group (F(1,10) = 57.9745, p < 0.001). However, the main effects of the recording device (F(2,20) = 1.5483, p = 0.2370) and experimental condition (F(1,10) = 1.2502, p = 0.2897) did not reach statistical significance. Likewise, the interactions between device and condition (F(2,20) = 0.2764, p = 0.7613) and between device, condition, and channel group (F(2,20) = 2.2954, p = 0.1266) were not significant. The only noteworthy interactions were observed between device and channel (F(2,20) = 4.6081, p < 0.05) and between condition and channel (F(1,10) = 15.6154, p < 0.05).

The beta power was notably higher for the posterior channels compared to the anterior channels (t = 13.142, p < 0.001). For the anterior channels, the beta power was more stronger during the eyes-open condition compared to the eyes-closed condition (t = 3.272, p < 0.05). Moreover, NVX recordings displayed greater beta power for the posterior channels compared to both Neiry Headband (t = 6.919, p < 0.001) and Muse (t = 3.243, p < 0.05).

The results from the ANOVA analysis for high beta power (Fig. 3, F) yielded several distinct findings. The main effects of the recording device (F(2,20) = 5.5221, p < 0.05), experimental condition (F(1,10) = 8.1142, p < 0.05), and channel group (F(1,10) = 22.1371, p < 0.001) were all found to be significant. However, the interactions between device and condition (F(2,20) = 1.6315, p = 0.2206) and between device, condition, and channel group (F(2,20) = 1.1531, p = 0.3358) were not significant. Additionally, the interactions between device and channel group (F(2,20) = 4.9652, p < 0.05) and between condition and channel group (F(1,10) = 18.0802, p < 0.05) were significant.

During the eyes-closed condition, high-beta power was significantly suppressed compared to the eyes-open condition (p = -5.6774, p < 0.001). Conversely, for comparison of the eyes-open condition versus eyes-closed condition, high-beta power was higher over the anterior sites (t = 5.903, p < 0.001). Moreover, high-beta power was higher over the posterior than anterior channels (p = 7.7817, p < 0.001).

In terms of the recording devices, the NVX recordings had a higher high-beta power compared to Muse recordings (t = 2.4525, p < 0.05) and Neiry Headband recordings (t = 6.1265, p < 0.001). Additionally, the Muse recordings had a elevated high-beta power compared to the Neiry Headband recordings (t = 3.137, p < 0.001).

Considering the posterior channels specifically, the high beta power was suppressed for the Neiry Headband recordings compared to NVX (t = -8.956, p < 0.001) and Muse (t = -3.795, p < 0.001) while it was elevated for the NVX compared to Muse (t = 4.036, p < 0.001).

The ANOVA analysis conducted for the gamma frequency range (Fig. 3, G) revealed significant main effects for device (F(2,20) = 7.4974, p < 0.05), condition (F(1,10) = 10.4112, p < 0.05), and channel group (F(1,10) = 16.4423, p < 0.05).

However, the interactions between device and condition (F(2,20) = 2.6102, p = 0.0984) and between device, condition, and channel group (F(2,20) = 1.5608, p = 0.2345) were not statistically significant. In contrast, the interactions between device and channel (F(2,20) = 4.0474, p < 0.05) as well as between condition and channel (F(1,10) = 11.7374, p < 0.05) were significant.

Similar to the the findings for high beta, the gamma power was lower during eyes closed than eyes open (t = -6.5174, p < 0.001). This effect was strongest for the posterior channels (t = 6.7493, p < 0.001). Yet, this effect was significant only for NVX (t = 6.426, p < 0.001) and Muse (t = 3.906, p < 0.001).

Comparing the recording devices, the gamma power recorded by Muse was lower compared to the NVX recordings (t = -3.7136, p < 0.05) recordngs and higher compared to Neiry Headband recordings (t = 3.2283, p < 0.05). Additionally, gamma power was lower for Neiry Headband as compared to NVX (t = -6.9752, p < 0.001).

Considering the signal spatial properties, the posterior channels of Neiry Headband had lower gamma power compared to both NVX (t = -9.310, p < 0.001) and Muse (t = -3.913, p < 0.001). Conversely, the posterior channels of NVX had higher gamma power compared to Muse (t = 4.595, p < 0.001).

#### 3.1.3. Correlation analysis

The correlation analysis of mean log PSD values revealed that across all frequency ranges, EEG signals of the Neiry Headband matched the conventional EEG recordings obtained with NVX better than those obtained with Muse. Additionally, the correlations between the NVX and Muse were weaker than the correlations observed between the Neiry Headband and NVX. (Fig. 4).

**Figure 4:**
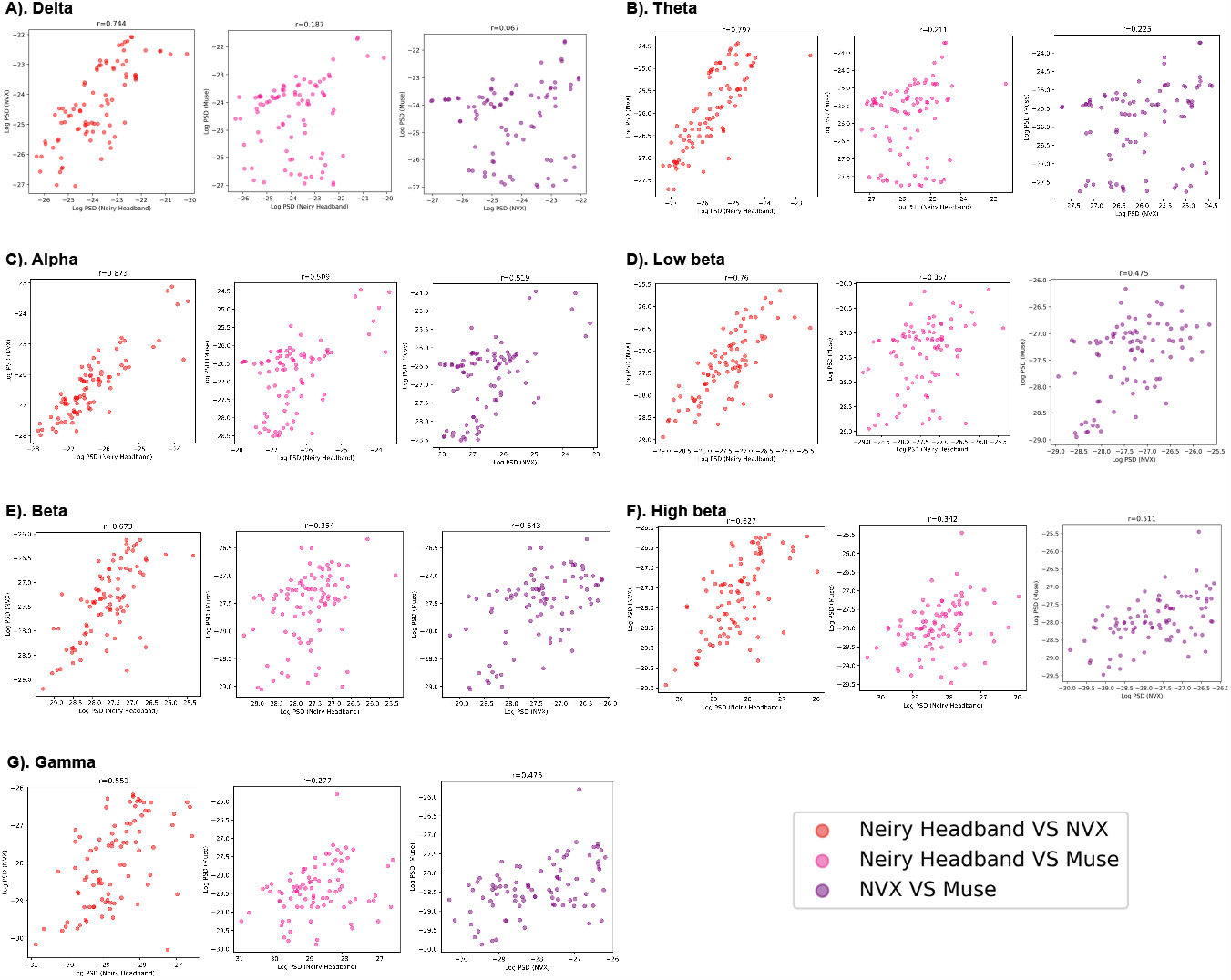
The correspondence between the recordings obtained with different devices (Neiry Headband, NVX and Muse) assessed as pairwise correlations of mean log PSD values for different frequency range

As it is clear from Table 1, most of the correlation values were found to be statistically significant with the exception for two values for the delta frequency band: the correlation between Neiry Headband and Muse (r(86)=0.187, p=0.081), and the correlation between NVX and Muse (r(86)=0.067, p=0.535). For all other frequency ranges, the correlations were significant. The correlation was the highes for the comparison of the alpha-range signals of the Neiry Headband and NVX (r(86)=0.873, p<0.001).

**Table 1:**
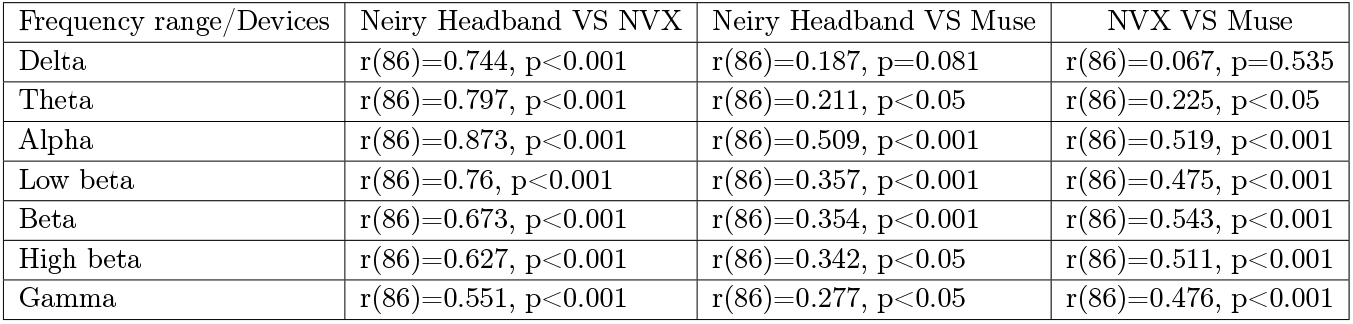
Results of correlation analysis for Neiry Headband, NVX and Muse.

### 3.2. Study 2. Neiry Headphones

#### 3.2.1. Resting-state EEG

In Figure 5, we present a 10-second sample of resting-state EEG data, illustrating both the eyes-open and eyes-closed conditions for Neiry Headphones, NVX, and Muse Headband. A visual inspection reveals the presence of alpha spindles in both Neiry Headphones and NVX when the eyes are closed. Furthermore, there is apparent evidence of low-frequency drifts in the sample from Neiry Headphones, when the eyes are open.

**Figure 5:**
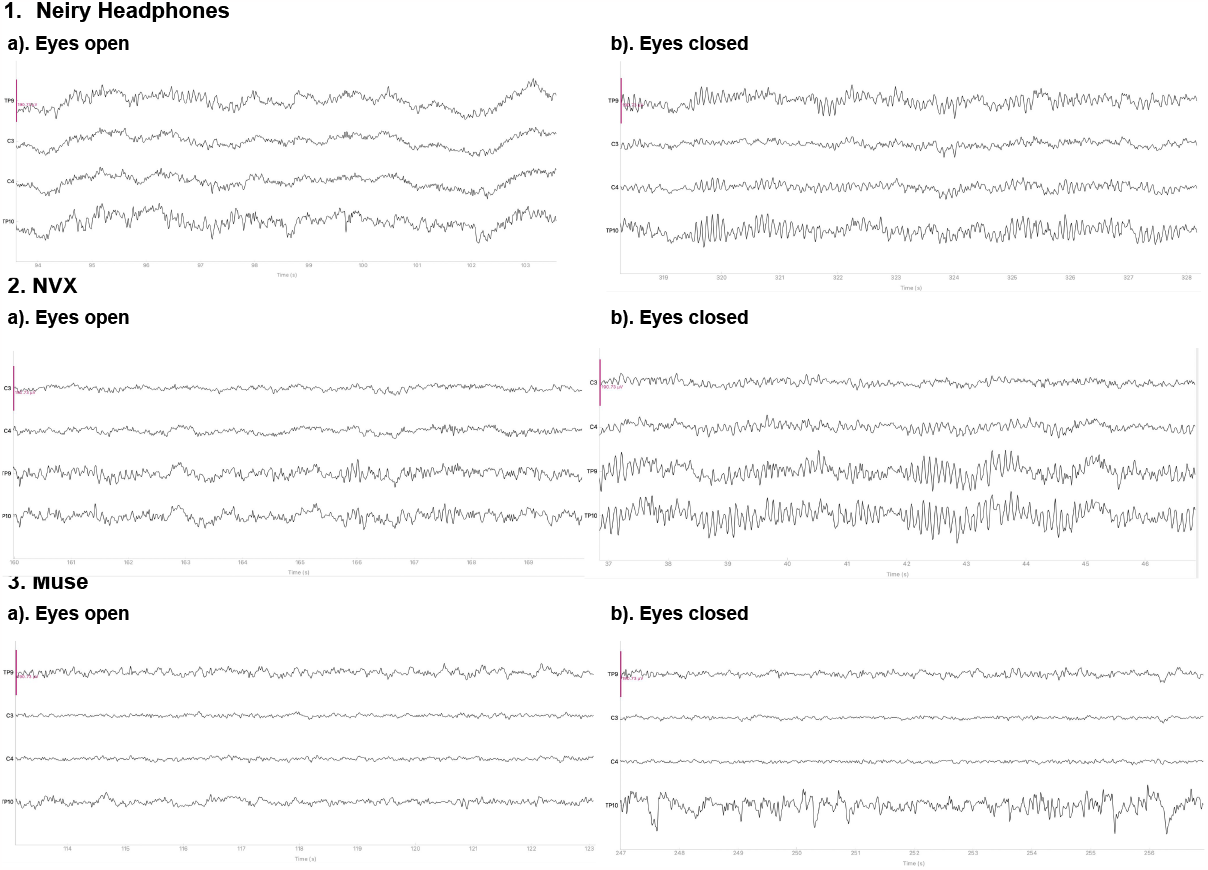
Representative samples of the recordings with the Neiry Headphones (1), NVX (2) and Muse (3) devices for the eyes-open and eyes-closed conditions. The y-limit is set to ±190.73 mV for all graphs. The signal was bandpass-filtered from 0.5 to 30 Hz for visualization.

**Figure 6:**
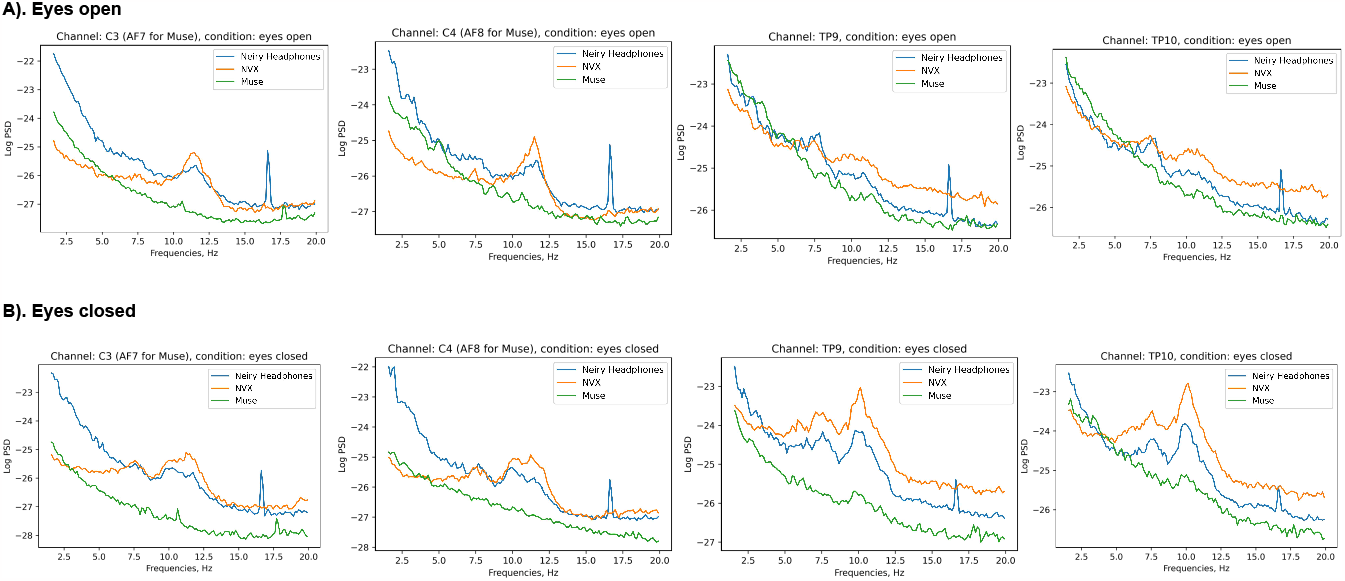
Across-subject (N=13) averages of log PSD demonstrating the difference in alpha power during eyes-open (A) versus eyes-closed (B) conditions for Neiry Headphones, NVX and Muse

#### 3.2.2. Spectral analysis

Figure 2 shows the across-participant average of log PSD, clearly showing an increase in alpha power when the eyes were closed, especially for NVX and Neiry Headphones devices.

In a manner akin to Study 1, we employed a repeated measures ANOVA to investigate the impact of device, condition, and channel group on the mean log PSD within a specified frequency range.

There was a significant main effect of the device on delta power (Fig. 7, A) (F(2, 24) = 7.684, p < 0.05). Likewise, there were main effects observed for condition (F(1, 12) = 11.665, p < 0.05) and channel group (F(1, 12) = 115.156, p < 0.001). Additionally, there were significant interactions between device and condition (F(2, 24) = 9.437, p < 0.001) and between device and channel group (F(2, 24) = 18.803, p < 0.001). Conversely, interactions between condition and channel group were not found to be significant (F(1, 12) = 0.117, p = 0.7385), nor were interactions between device, condition, and channel group (F(2, 24) = 0.019, p = 0.982).

**Figure 7:**
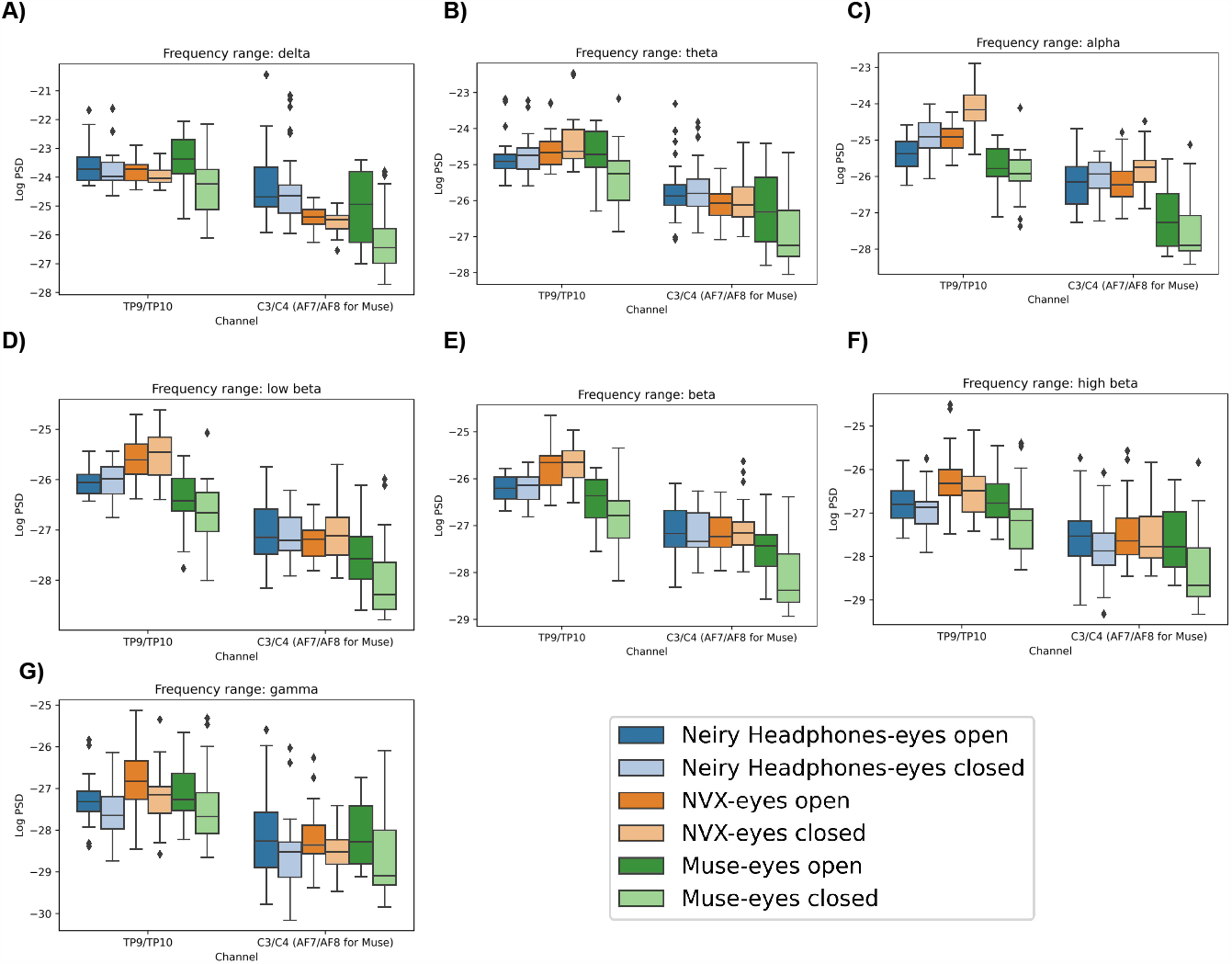
Boxplots showing the distribution of mean log PSD values across conditions and channel groups for different frequency ranges: delta (A), theta (B), alpha(C), low beta (D), beta (E), high beta (F), gamma (G) for Neiry Headphones, NVX and Muse

Subsequent post-hoc analysis illuminated specific insights. Delta power was significantly lower in both the Muse (t = -5.368, p < 0.001) and NVX (t = -5.9088, p < 0.001) devices compared to Neiry Headphones. Furthermore, in the eyes-open condition, delta power demonstrated a significant increase (t = 5.4733, p < 0.001). This increase was observed specifically for NVX (t=4.965, p<0.001) and Muse (t=5.842, p<0.001).

Notably, for anterior channels (C3 and C4 for Neiry Headphones and NVX, AF7 and AF8 for Muse), delta power was significantly lower (t = -14.7414, p < 0.001) than for posterior channels (TP9 and TP10).

Delta power in Neiry Headphones was significantly higher than in NVX, both in eyes-open (t = 3.889, p < 0.001) and eyes-closed (t = 4.439, p < 0.001) conditions. Moreover, it was higher than in Muse, but only in the eyes-closed condition (t = 5.910, p < 0.001).

Specifically, delta power in Neiry Headphones at C3 and C4 channels was significantly higher than in NVX (t = 6.456, p < 0.001) and Muse (AF7 and AF8 channels) (t = 6.180, p < 0.001).

In the theta band (Fig. 7, B), significant effects were observed for device (F(2,24) = 7.135, p < 0.05) and channel group (F(1,12) = 263.663, p < 0.001). However, the main effect of condition was not significant (F(1,12) = 2.038, p = 0.1789). Notably, there were significant interactions between device and condition (F(2,24) = 11.363, p < 0.001) as well as between device and channel group (F(2,24) = 12.099, p < 0.001). Conversely, the interactions between condition and channel were not significant (F(1,12) = 0.069, p = 0.7974), and the interactions involving device, condition, and channel group were also not significant (F(2,24) = 0.2169, p = 0.8066).

Post-hoc analysis revealed that theta power exhibited significant reductions when comparing Muse Headband to both NVX (t = -4.951, p < 0.001) and Neiry Headphones (t = -5.729, p < 0.001). Furthermore, theta power was significantly lower in anterior channels (C3 and C4 for Neiry Headphones and NVX, and AF7 and AF8 for Muse) compared to posterior channels (TP9 and TP10) (t = -21.4988, p < 0.001).

When examining the impact of the condition, it was found that theta power significantly decreased in the eyes closed condition with the Muse Headband compared to Neiry Headphones (t = -6.845, p < 0.001) and NVX (t = -6.559, p < 0.001). Additionally, NVX exhibited a significant increase in theta power in the eyes closed condition (t = 4.328, p < 0.001), while Muse showed a decrease in theta power during the eyes-closed condition (t = -4.736, p < 0.001).

Furthermore, Neiry Headphones demonstrated an increase in theta power at C3 and C4 channels compared to NVX (t = 6.456, p < 0.001) and Muse (AF7 and AF8 channels) (t = 6.180, p < 0.001).

In the alpha band analysis (Fig. 7, C), we observed the following significant effects. Firstly, there was a significant main effect of the device (F(2,24)=89.064, p<0.001) and a significant main effect of the channel (F(1,12)=242.938, p<0.001). Although the main effect of the condition did not reach significance (F(1,12)=4.475, p=0.056), there was a noticeable trend. Furthermore, we found significant interactions between device and condition (F(2,24)=12.135, p<0.001), device and channel group (F(2,24)=10.002, p<0.001), and condition and channel group (F(1,12)=7.0962, p<0.05). However, the interaction between device, condition, and channel group was not significant (F(2,24)=0.4, p=0.6746).

Delving into post-hoc analysis, we observed that alpha power significantly increased during eyes-closed conditions at the posterior channels TP9 and TP10 (t=4.601, p<0.001). Additionally, alpha power was significantly reduced in Muse compared to NVX (t=-15.057, p<0.001) and Neiry Headphones (t=-12.004, p<0.001), while NVX exhibited significantly higher alpha power compared to Neiry Headphones (t=7.2804, p<0.001).

Notably, in the anterior channels (C3 and C4 for Neiry Headphones and NVX, AF7 and AF8 for Muse), alpha power was significantly suppressed (t=-24.455, p<0.001) compared to the posterior channels (TP9 and TP10).

Furthermore, the significant increase in alpha power due to eyes closed conditions was only observed in Neiry Headphones (t=4.571, p<0.001) and NVX (t=8.85, p<0.001).

Lastly, alpha power was more pronounced at anterior channels (C3 and C4) in Neiry Headphones (t=10.434, p<0.001) and NVX (t=9.916, p<0.001) compared to Muse (AF7 and AF8 channels). Additionally, at posterior channels TP9 and TP10, alpha power in Neiry Headphones was reduced compared to NVX (t=-12.185, p<0.001), while Muse exhibited reduced alpha power at those channels compared to Neiry Headphones (t=-7.134, p<0.001) and NVX (t=11.508, p<0.001).

In the low beta frequency range (Fig. 7, D), we observed significant effects across several factors. There was a significant main effect of the device used (F(2,24)=27.284, p<0.001). Likewise, there was a significant main effect of the channel group (F(1,12)=222.06, p<0.001). However, the main effect of the condition was not statistically significant (F(1,12)=1.64, p=0.2245). Furthermore, we observed significant interactions between the device and condition (F(2,24)=5.328, p<0.05) as well as between the device and channel group (F(2,24)=6.245, p<0.05). However, interactions between the condition and channel group were not significant (F(1,12)=0.3408, p=0.5702), nor were interactions between the device, condition, and channel group (F(2,24)=1.275, p=0.2977).

Post-hoc analysis revealed specific differences in low beta power. Muse exhibited reduced low beta power compared to both NVX (t=-10.267, p<0.001) and Neiry Headphones (t=-8.215, p<0.001). Neiry Headphones also displayed suppressed low beta power compared to NVX (t=-3.9355, p<0.001), particularly in the eyes closed condition (t=-4.121, p<0.001) and in posterior channels (t=-9.057, p<0.001). Additionally, low beta power was consistently lower in anterior channels (C3 and C4 for NVX and Neiry Headphones, AF7 and AF8 for Muse) compared to posterior channels (TP9 and TP10) (t=-26.27, p<0.001). Muse, in particular, exhibited increased low beta power when the eyes were open (t=3.788, p<0.05).

In the beta frequency range (Fig. 7, E), similar statistical patterns emerged. There was a significant main effect of the device (F(2,24)=23.755, p<0.001), a main effect of the condition (F(1,12)=5.663, p<0.05), and a main effect of the channel group (F(1,12)=220.248, p<0.001) on beta power. Additionally, we found significant interactions between the device and condition (F(2,24)=7.546, p<0.05) and between the device and channel group (F(2,24)=5.611, p<0.05). However, interactions between the condition and channel group (F(1,12)=0.753, p=0.4025) and between the device, condition, and channel group (F(2,24)=0.6117, p=0.5507) were not statistically significant.

Post-hoc analysis in the beta frequency range indicated that Muse had reduced beta power compared to both NVX (t=-10.364, p<0.001) and Neiry Headphones (t=-7.551, p<0.001). Neiry Headphones also exhibited suppressed beta power compared to NVX (t=-4.792, p<0.001), particularly in the posterior channels TP9 and TP10 (t=-9.663, p<0.001). Moreover, beta power was significantly higher in the eyes open condition compared to the eyes closed condition (t=3.411, p<0.001). Similar to the low beta range, beta power was consistently lower in anterior channels (C3 and C4 for NVX and Neiry Headphones, AF7 and AF8 for Muse) compared to posterior channels (TP9 and TP10) (t=-25.275, p<0.001). Muse also exhibited increased beta power when the eyes were open (t=4.909, p<0.001).

In the high beta frequency range (Fig. 7, F), we observed significant main effects for device (F(2,24)=5.5835, p<0.05), condition (F(1,12)=26.327, p<0.001), and channel group (F(1,12)=90.719, p<0.001). Notably, the only significant interaction was between the device and condition (F(2,24)= 4.444, p<0.05), while interactions between device and channel group (F(2,24)=1.666, p<0.2101), between condition and channel group (F(1,12)=1.412, p=0.2578), and between device, condition, and channel group (F(2,24)=2.272, p=0.1249) were not statistically significant.

Post-hoc analysis indicated that NVX had higher high beta power compared to Muse (t=6.255, p<0.001) and Neiry Headphones (t=6.326, p<0.001). In the eyes open condition, high beta power was more pronounced than in the eyes closed condition (t=7.148, p<0.001). Additionally, in the eyes closed condition, Neiry Headphones exhibited higher high beta power than Muse (t=3.16, p<0.05). As with previous frequency ranges, high beta power was consistently lower in anterior channels (C3 and C4 for NVX and Neiry Headphones, AF7 and AF8 for Muse) compared to posterior channels (TP9 and TP10) (t=-18.883, p<0.001).

In the gamma frequency range (Fig. 7, G), we found no significant main effect of the device (F(2,24)=0.617, p=0.5478). However, we did observe significant main effects for both the condition (F(1,12)=45.948, p<0.001) and the channel group (F(1,12)=75.7, p<0.001). There were no significant interactions between device and condition (F(2,24)=0.49, p=0.6189), between device and channel group (F(2,24)=1.7622, p=0.1932), between condition and channel group (F(1,12)=0.816, p=0.3841), or among device, condition, and channel group (F(2,24)=1.572, p=0.2283). In the eyes open condition, gamma power was significantly higher than in the eyes closed condition (t=8.121, p<0.001). Additionally, gamma power was significantly lower in anterior channels (C3 and C4 for NVX and Neiry Headphones, AF7 and AF8 for Muse) compared to posterior channels (TP9 and TP10) (t=-17.382, p<0.001).

#### 3.2.2. Correlation analysis

The examination of mean log PSD values through correlation analysis showed that, in all frequency ranges, except delta band, the EEG signals from Neiry Headphones closely aligned with conventional EEG recordings from NVX, surpassing those from Muse (see Fig. 8).

**Figure 8:**
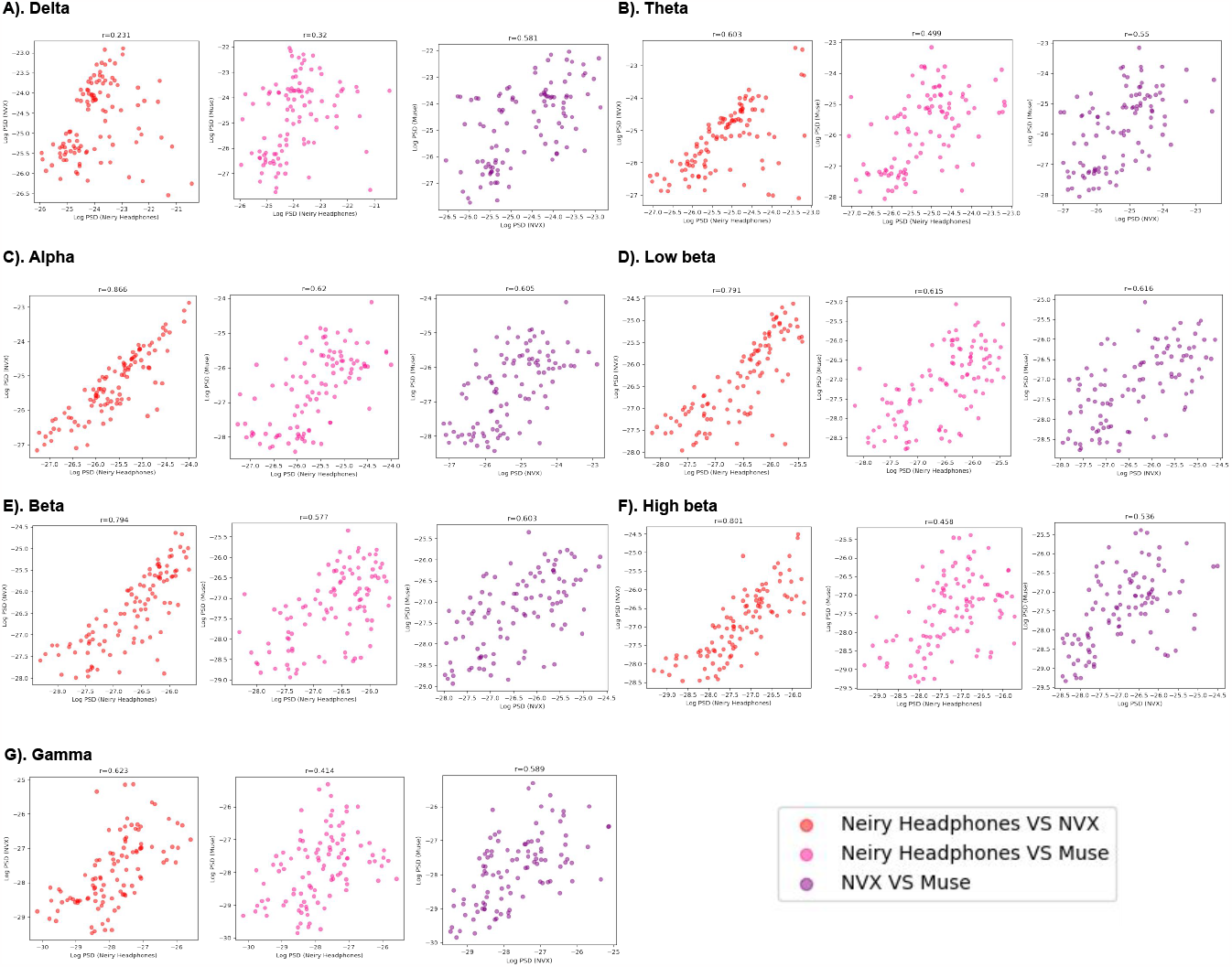
The correspondence between the recordings obtained with different devices (Neiry Headphones, NVX and Muse) assessed as pairwise correlations of mean log PSD values for different frequency ranges

As evident from Table 2, all the correlation values were statistically significant. Notably, the alpha band exhibited the highest correlation between Neiry Headphones and NVX (r(102)=0.866, p<0.001). For the low beta, beta, and high beta ranges, the correlation between Neiry Headphones and NVX surpassed that between Muse and NVX. In both the theta and gamma ranges, the correlation values were comparable. In the delta range, the correlation between Neiry Headband and NVX was considerably lower than that between NVX and Muse (r(102)=0.231, p<0.05 againstr (102)=0.581, p<0.001).

**Table 2:** Results of correlation analysis for Neiry Headphones, NVX and Muse.

| Frequency range/Devices | Neiry Headphones VS NVX | Neiry Headband VS Muse | NVX VS Muse |
| --- | --- | --- | --- |
| Delta | $r(102)=0.231, p<0.05$ | $r(102)=0.32, p<0.05$ | $r(102)=0.581, p<0.001$ |
| Theta | $r(102)=0.603, p<0.001$ | $r(102)=0.499, p<0.001$ | $r(102)=0.55, p<0.001$ |
| Alpha | $r(102)=0.866, p<0.001$ | $r(102)=0.62, p<0.001$ | $r(102)=0.605, p<0.001$ |
| Low beta | $r(102)=0.791, p<0.001$ | $r(102)=0.615, p<0.001$ | $r(102)=0.616, p<0.001$ |
| Beta | $r(102)=0.794, p<0.001$ | $r(102)=0.577, p<0.001$ | $r(102)=0.603, p<0.001$ |
| High beta | $r(102)=0.801, p<0.001$ | $r(102)=0.458, p<0.001$ | $r(102)=0.536, p<0.001$ |
| Gamma | $r(102)=0.623, p<0.001$ | $r(102)=0.414, p<0.001$ | $r(102)=0.589, p<0.001$ |

## 4. Discussion

In the present study, we obtained resting-state EEG data using several recording devices. Although unsophisticated, monitoring resting-state cortical activity has multiple medical and consumer applications, so perfecting these kind of recordings and making them affordable and easy to implement offers multiple benefits. Yet, the potential users of EEG equipment would also benefit from a realistic assessment of the performance of these systems. Here we compared the recordings obtained with three consumer-grade EEG devices, the Neiry Headband, the Neiry Headphones and the Muse Headband, with the data collected with the medical-grade device, the NVX. We found distinct sets of signal-quality parameters that could be applied to the comparison of different recording devices, illuminating their potential advantages and limitations for various EEG-based applications.

The performance in the delta-rhythm band is of great interest because of the applications where awake and sleep (not considered here) states could be assessed. Here, the Neiry Headband consistently had a higher delta power than the other devices. This was particularly clear for the eyes open condition, and for the T3/T4 channels of Neiry. By contrast, the AF7/AF8 of Muse had an attenuated delta power. While this finding suggests that the Neiry Headband is suitable for the monitoring of low-frequency EEG components (Hinrichs et al., 2020), this frequency range is also prone to low-frequency artifacts which are difficult to control in consumer settings. Yet, we should not dismiss the low-frequency signal as merely noisy particularly because the Neiry Headset was capable of detecting an increase in delta power (as well as theta power) during eyes-open conditions, which aligns with the idea of unstructured processing of the environment (Barry et al., 2007). The same effect was seen in the NVX recordings, which supports the validity of the Neiry Headband’s findings. Accordingly, our tests lend credibility to the Neiry Headband’s data in the low-frequency range. In Study 1, the Muse recordings detected this modulation only at the posterior channels, which questions the reliability of delta-rhythm monitoring recordings with this device.

In contrast, Neiry Headphones exhibited a greater susceptibility to low-frequency artifacts. The delta power in recordings from this device exceeded that of Muse and NVX. Unlike Muse and NVX, this delta power was not modulated by eye conditions.

For the theta-frequency range, our results showed that the Neiry Headband exhibited suppressed theta power compared to Muse for the posterior channels, while it yielded higher theta power than Muse for the anterior channels. The NVX system also exhibited unique patterns. In particular, its posterior theta power (O1 and O2) was lower compared to Muse and its anterior theta power (T3 and T4) was higher compared to Neiry Headband. These spatial difference could become important for the applications based on the comparison of theta power in different cortical regions. Notably the Neiry Headband and NVX successfully detected increased theta power when the eyes were closed, which could be considered a standard spectral change (Barry et al., 2007). On the contrary, while NVX still recorded a notable rise in theta power at channels C3, C4, TP9 and TP10, theta power remained unchanged in Neiry Headphones regardless of whether the eyes were open or closed.

The alpha frequency range is perhaps the most reliable feature of these kind of recordings because alpha oscillations are typically very prominent and could be easily distinguished from the artifacts. In this range the Muse device was less sensitive compared to the NVX and Neiry Headband recordings, which, combined with the consideration of the Muse reduced spatial coverage, suggests this device does not effectively capture alpha activity, particularly over the posterior brain regions. By contrast, the posterior-alpha rhythm was reliably detected by the Neiry Headband and NVX. In particular, these devices detected posterior alpha suppression when the eyes were open (Barry et al., 2007). When the eyes were closed and posterior alpha activity was highly synchronized, the Neiry Headband and the NVX system signals were stronger compared to the Muse Headband. Similar findings were observed with Neiry Headphones, where an increase in alpha power occurred when participants closed their eyes, mirroring the results seen with NVX. However, it’s noteworthy that the overall alpha power in Neiry Headphones was lower than that in NVX.

As to the low-beta, beta, and high-beta and gamma activity over the posterior regions, the NVX was consistently more sensitive compared to both Neiry Headband and Muse recordings in Study 1. In the second study, Neiry Headphones exhibited a notable suppression of low beta, beta, and high beta activity compared to NVX, particularly at posterior channels. Moreover, all three devices demonstrated increase in beta power, when eyes were open. Intriguingly, the state of the eyes influenced gamma power in NVX and Muse, suppressing it when the eyes were closed; however, this effect was not observed with Neiry Headphones.

he correlation analysis indicates that, across all frequency ranges, the Neiry Headband exhibited a closer match with NVX readings compared to Muse. Notably, the correlations between Neiry Headphones and NVX were higher than those between Neiry Headband and NVX in the low beta, beta, high beta, and gamma ranges. This phenomenon could be attributed to the Neiry Headband’s electrode placements being more sensitive to detecting high-frequency oscillations. Generally, Niery Headphones displayed stronger correlations with NVX than Muse did in the theta, alpha, low beta, beta, and high beta ranges. However, in the delta band, the correlation between Neiry Headphones and NVX was notably low, and in the theta range, it was lower than the correlation between Neiry Headband and NVX. Nevertheless, for the alpha range, the correlations were similar between Neiry Headphones and NVX as well as between Neiry Headband and NVX.

We conclude that Neiry’s dry-electrode technology shows an overall consistency with the standard medical-grade measurements.

Overall, our results suggest that such dry-electrode systems as Neiry Headband with channels at T3, T4, O1 and O2 are appropriate for such consumer application as cognitive monitoring where delta, theta and alpha frequency readings are used to assess the cognitive state. Systems like Neiry Headphones, featuring channels at C3, C4, TP9, and TP10, tend to exhibit a higher susceptibility to low-frequency artifacts. They may not be as responsive to low-frequency modulations within the theta range, but they are capable of capturing activities within the alpha range. Additionally, they can adequately represent modulations in beta power. Yet, when selecting the system to use, it is essential to consider the specific research objectives and frequency bands of interest. Thus, for tasks that involve high-frequency oscillations (beta and gamma), the gel electrode-based NVX system is still a more suitable choice due to its higher power sensitivity in these frequency ranges. One limitation of the study is the specific electrode configurations used for each device. Different electrode placements and spatial coverage could have contributed to the observed differences in signal quality.

In conclusion, our findings highlight the distinct characteristics of signal quality across different recording devices and frequency ranges. The results suggest that the choice of device significantly influences the measurements of EEG power in different frequency bands and brain regions. Researchers and practitioners should carefully consider the specific application and frequency ranges of interest when selecting and recommending an appropriate recording device. The modern dry electrode technology showed promising results, but its performance should be additionally evaluated based on the specific research context and experimental modulation of such cognitive states as relaxation, stress, mental workload, and the others.

Overall, this study contributes to the growing body of research on the dry electrode technology, its potential applications in EEG research and real-world EEG-based applications, and the development of unified requirements for consumer-grade applications of this type.

